# MSMICA: computational metabolite identification in untargeted metabolomics by integrating MS, retention time, and biological evidence

**DOI:** 10.64898/2026.08.15.744986

**Authors:** Jiada Zhan, Jaclyn Weinberg, William J. Crandall, Zhaohui Qin, Zachery R. Jarrell, Joshua D. Preston, Mary Nellis, Sami Teeny, Donghai Liang, Greg S. Martin, Nathan L. Price, Rafael de Cabo, Viraj Master, Barbara A. Cohn, Young-Mi Go, Dean P. Jones

**Affiliations:** Division of Pulmonary, Allergy, Critical Care, and Sleep Medicine, School of Medicine, Emory University, Atlanta, GA, USA; Nutrition and Health Sciences, Laney Graduate School, Emory University, Atlanta, GA, USA; Department of Biostatistics and Bioinformatics, Emory University, Atlanta, GA, USA; Medical Scientist Training Program, School of Medicine, Emory University, Atlanta, GA, USA; Translational Gerontology Branch, National Institute on Aging, National Institutes of Health, Baltimore, MD, USA; Gangarosa Department of Environmental Health, Rollins School of Public Health, Emory University, Atlanta, GA, USA; Division of Urology, School of Medicine, Emory University, Atlanta, GA, USA; Child Health and Development Studies, Public Health Institute, Oakland, CA, USA

## Abstract

Mass Spectrometry Metabolomics Identification Connection Algorithm (MSMICA) is an algorithm for automated metabolite identification in untargeted liquid chromatography-high-resolution mass spectrometry (LC-HRMS) analyses. Limitations in metabolite identification can occur due to the availability and cost of standards and prevent recognition of metabolic factors impacting human health and disease. MSMICA performs mass-to-charge-ratio matching with chemical structures and clusters of LC-HRMS features for adduct and isotope forms. A local optimization is then used to integrate retention time prediction, metabolite precursor-product and transporter correlations, and biospecimen-specific abundance information for metabolite identification. Applying MSMICA to various internal and external mammalian datasets, validation results showed a 96.2 ± 5.1% correct rate of metabolite identification. When multiple LC-HRMS datasets were used, MSMICA enabled greater metabolite identifications, expanded metabolic pathway coverage, and data harmonization. Thus, MSMICA applies multiple pieces of evidence to substantially improve metabolite identification coverage and accuracy for known metabolites.

## Introduction

Metabolomics links the genome, exposome, and health phenotypes in systems-level physiology and disease^1–3^. Liquid chromatography-high-resolution mass spectrometry (LC-HRMS) has emerged as the central technology for metabolomics, and extensive LC-HRMS datasets for humans and model systems are available in data repositories^4,5^. Unfortunately, more than half of the mass spectral features associated with human diseases and exposures are unidentified^6^. Even with accurate mass-to-charge ratios (*m/z*) obtained by HRMS, high rates of false-positive discoveries and redundancy can occur due to the untargeted nature of HRMS^7^. Current metabolomics identification consequently relies upon *m/z*, retention time (RT), and ion dissociation (MS/MS) spectral matching to chemical standards^8,9^. Practical limitations in metabolite identification arise from difficulties in running standards and from suboptimal MS/MS for low-intensity features. As a result, using untargeted metabolomics to build metabolic models is challenging, and metabolite identification remains a bottleneck in LC-HRMS-based untargeted metabolomics.

Existing computational approaches have been proposed to overcome this bottleneck, including tools related to metabolic pathway analysis^10–12^, multi-layer network analysis^13–18^, machine learning^19–21^, in-source fragmentation^22,23^, and Bayesian statistics^24–27^. Particularly, Mummichog offers a novel approach to predict biological pathway changes without upfront metabolite identifications^10^. NetID presents a global network optimization approach for metabolite discovery using adduction, fragmentation, isotope, and feasible biochemical transformations^17^. KGMN (MetDNA2) uses a knowledge-guided multi-layer network of metabolic reactions, MS/MS similarity, and global feature correlation for global metabolite annotation^16^. MetDNA3 applies a data-driven and knowledge-driven network algorithm to enhance the accuracy and coverage of metabolite annotation^18^. xMSannotator employs a multistage clustering algorithm for metabolite annotation using isotope/adduct patterns^14^. Nevertheless, all of the methods still face fundamental tradeoffs: accuracy, identity redundancy, and coverage, as many features have several identities in the output results, even though experimental RT, MS/MS, and quantification can confidently assign one identity to one feature in many cases. Moreover, MS/MS data is often mandatory to run many computational metabolomics annotation algorithms, and the quality of MS/MS data can become limiting for high-throughput analyses of low abundance metabolites.

In principle, biological evidence can improve the interpretation of untargeted metabolomics data because the metabolic precursor-product relationships, metabolic co-transporter relationships, and metabolite concentration in human biospecimens are well-documented in the literature and metabolite databases. Precursor-product relationships are critical in the identification of xenobiotic metabolites^28^. In addition, known metabolic precursor-product and co-transporter relationships have facilitated the identification of endogenous metabolites^29^. Theoretically, reported concentrations of metabolites in human biospecimens are also an important consideration for metabolite identification because they reflect the steady-state condition of mammalian metabolism, which may help distinguish metabolites with the same monoisotopic mass by known abundance. Because biological evidence is independent of mass spectrometry evidence (mass accuracy) and liquid chromatography evidence (RT), integrating them into a computational interpretation of untargeted metabolomics data may be well-suited for obtaining metabolic network information and associated metabolite identifications.

In this study, we develop a computational tool, MSMICA (<u>M</u>ass <u>S</u>pectrometry <u>M</u>etabolomics Identification <u>C</u>onnection <u>A</u>lgorithm), to integrate metabolic network information from LC-HRMS untargeted metabolomics with associated metabolite identifications, without relying on MS/MS collection or in-house RT libraries. MSMICA takes a feature intensity table containing *m/z*, RT, and integrated ion intensities across samples as input, and assigns metabolite identities to features by combining multiple lines of evidence, including accurate-mass matching, adduct and isotopic clustering, RT prediction, precursor-product or shared-transporter correlations, and the alignment of feature intensity with estimated metabolite concentration. This workflow is designed to reduce ambiguity in metabolite identification and provide streamlined interpretation of metabolite information and identification. MSMICA offers an automated, transferable assessment for metabolite identification probability across different LC-HRMS untargeted metabolomics datasets^30^ and can be used to harmonize metabolomics data from different experimental settings.

## Results

### MSMICA algorithm

MSMICA has four computational steps: *m/z* matching, parameter estimation, clustering of adducts and isotopes, and local optimization per monoisotopic mass.

As a prerequisite, MSMICA depends on the feature table generated from the raw LC-HRMS data. It uses the feature table for metabolite *m/z* matching to a metabolite database (**Fig. 1 and Supplementary Table 1**). The metabolite database was created by merging HMDB and KEGG metabolite databases (**Supplementary Data 1**)^31,32^. MSMICA annotates MS^1^ features with user-specified adduct forms of metabolites based on accurate mass, with a default tolerance of ± 10 ppm. Chemically plausible adducts are determined from metabolite structure and classification (**Supplementary Note 1)**.

**Figure 1.**
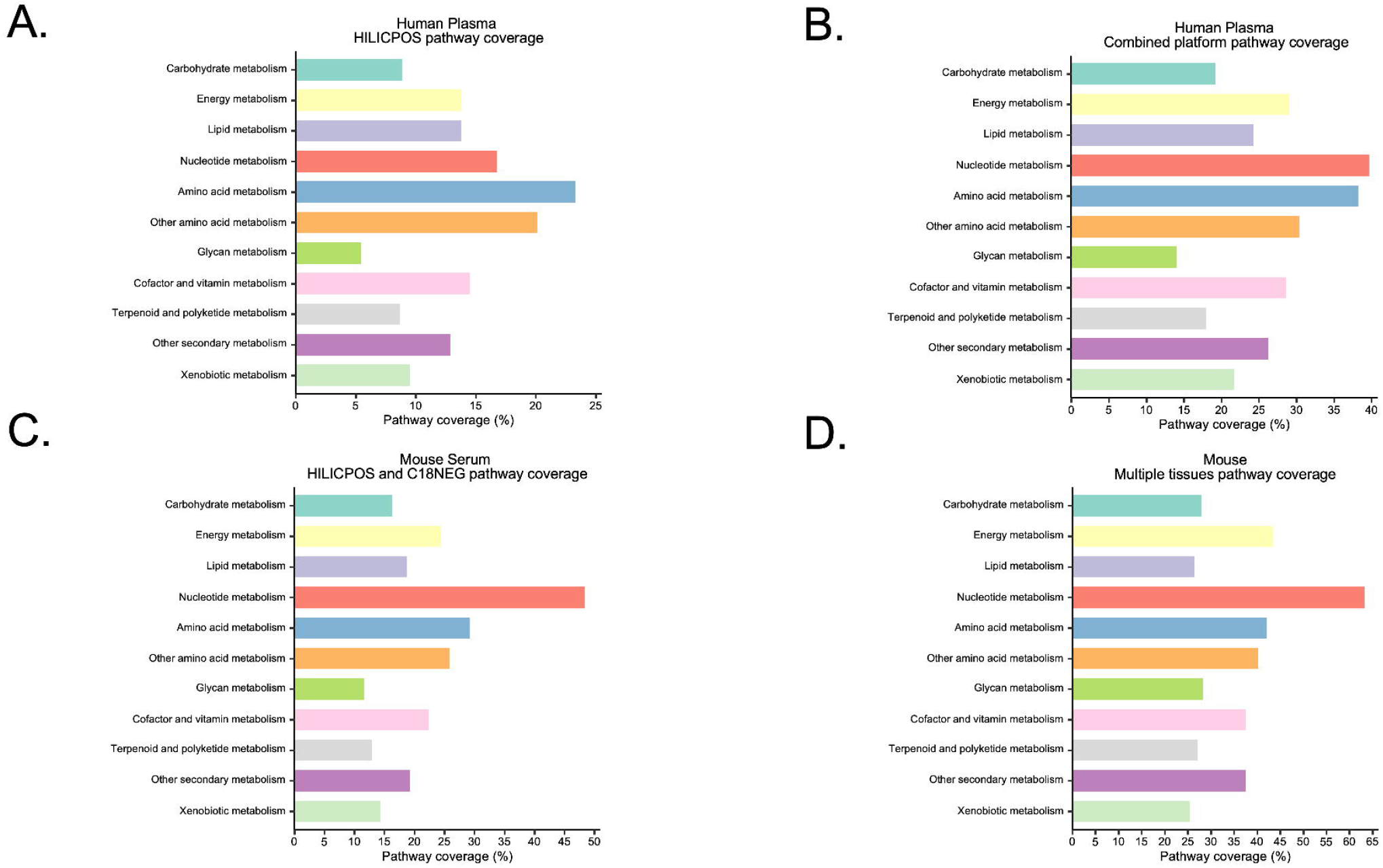
Overview of MSMICA. The input data is a feature table with the *m/z* as the first column, RT as the second column, and integrated ion intensities (peak areas) of samples as the remaining columns. MSMICA’s output is a list of features with the most likely metabolites based on the probability generated by MSMICA and their adduct forms selected. In most cases, one feature is assigned to one identity. MSMICA selects the most likely features for each metabolite using *m/z* matching with a chemical structure check, clustering of adducts and isotopes, and local optimization for metabolites with the same monoisotopic mass via RT prediction, precursor-product/transporter correlations, and feature intensity metabolite concentration alignment.

Next, MSMICA uses unique *m/z* annotated metabolites with non-overlapping isomers in chromatography as anchor metabolites for RT mapping. These anchor metabolites are used to align RTs with the RT reference library by a monotonically constrained generalized additive model (GAM). The fitted GAM is then used to predict the RTs of other metabolites. Using the same anchor metabolites, MSMICA also estimates model parameters for clustering of adducts and isotopes (RT difference and correlation thresholds), RT prediction error, and enzymatic precursor-product and shared-transporter correlations, hereafter referred to as precursor-product/transporter correlations. Specifically, this step estimates the standard deviation of the RT prediction error, along with the mean and standard deviation of the precursor-product/transporter correlations. These parameters are used in later steps of the algorithm.

MSMICA then applies a clustering algorithm to remove redundant features derived from the same metabolite (**Fig. 1**). This step identifies features that share the same mass but arise from different isotope forms (^12^C, ^13^C) and adduct forms (e.g., +H^+^, +Na^+^). MSMICA then uses isotope and adduct correlation patterns within a Bayesian framework to select the features most likely representing a single metabolite based on the predicted mass. Only chemically plausible major adducts are retained for later analysis; features corresponding to other adducts are excluded.

After redundant features are excluded, MSMICA performs local optimization for metabolite identification within each monoisotopic mass group. For each feature-metabolite pair, the default setting of MSMICA estimates the probability of a correct assignment using 3 sources of evidence: RT prediction error, average precursor-product/transporter correlation, and feature intensity metabolite concentration alignment. The feature intensity metabolite concentration alignment favors the feature-metabolite pair in which relatively high intensity features are paired with relatively abundant metabolites in the algorithm’s specified biospecimen (default is blood). All pieces of evidence are used to calculate a log score for each feature-metabolite pair. These scores are then converted to posterior probabilities by softmax normalization. Then, starting at the most abundant metabolite, the feature-metabolite pair with the highest posterior probability is selected first, and other pairs are removed. After each iteration, selected feature-metabolite pairs are removed from the candidate set. The procedure continues for the next most abundant metabolite until no new feature-metabolite pairs can be assigned. Finally, MSMICA classifies results by Schymanski identification Levels 3a, 3b, and 4 according to the number and type of evidence supporting each metabolite identification (**Supplementary Table 2**).

### MSMICA demonstration and validation

To illustrate how MSMICA works, we use two features associated with a monoisotopic mass of 131.0582 Da. MSMICA first applies its clustering algorithm and detects multiple coeluting, correlated features around an RT of 80 s (**Fig. 2A**). This step removes redundant features from later analysis and supports that the feature at *m/z* 132.0656 and 80 s (feature 1) is very likely to correspond to a metabolite with neutral mass 131.0582 Da.

**Figure 2.**
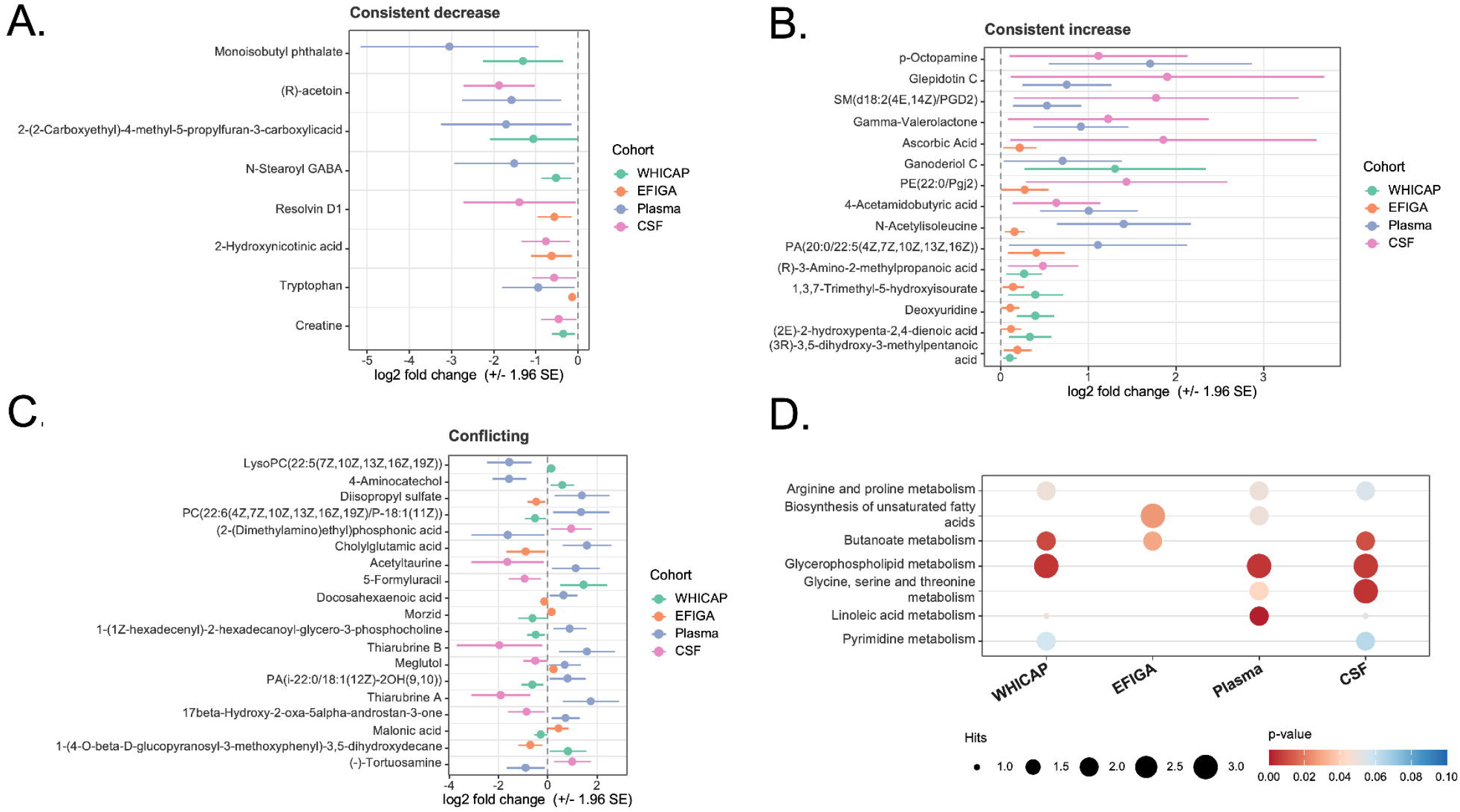
MSMICA demonstration and validation. **(A).** Example of finding multiple features with the monoisotopic mass of 131.0582 Da based upon RT coelution and correlation of adducts and isotopes in one human plasma LC-HRMS study. This is an example of Step 3 of the MSMICA algorithm: clustering of adducts and isotopes. The left panel contains the chromatograms for features with different adduct and isotopic forms with the exact mass in a plasma sample. Right panel describes the correlations between the features across all plasma samples. **(B)**. Example of using the local optimization per monoisotopic mass workflow to identify the feature with *m/z* 132.0656 and RT 80 s as hydroxyproline and the feature with *m/z* 132.0655 and RT 36 s as acetylalanine. The top panel is the data input of the local optimization algorithm with estimated concentration in blood, mean intensity, predicted retention time, actual retention time, and correlation with its proposed precursors and products among existing annotated metabolites. The bottom panel is the data output of the algorithm that demonstrates how the previous *m/z* -based metabolite annotations became computational metabolite identifications by considering mass spectrometry, liquid chromatography, and biological evidence. **(C)**. Validation of the MSMICA metabolite identifications by using RT and MS/MS of authentic standards. The left panel shows the chromatogram and RT when the standards were added or not. Right panel describes the MS/MS comparisons of the measured features with computational metabolite identifications and chemical standards. **(D)**. Summary of the fraction correct rate of MSMICA results in 4 independent plasma LC-HRMS studies and 3 serum LC-HRMS studies using Schymanski Level 1 identified metabolites (confirmed by 10 ppm *m/z* matching, RT within 30 s, MS/MS, and quantification with a check on HMDB for estimated blood concentration as the reference.

MSMICA then uses local optimization to assign metabolite identities to feature 1 and to a second feature at *m/z* 132.0655 and 36 s (feature 2). Both features match hydroxyproline and acetylalanine within the *m/z* tolerance, so all four feature-metabolite pairs are initially plausible (**Fig. 2B**). Based on the hypothesized metabolite identities, for each candidate pair, MSMICA evaluates predicted RT, the median precursor-product and transporter correlation coefficient, and feature intensity metabolite concentration alignment. Using this combined evidence, as hydroxyproline is the most abundant metabolite and acetylalanine is the second most abundant metabolite in blood, MSMICA first assigns feature 1 to hydroxyproline and then assigns feature 2 to acetylalanine in the next iteration of the local optimization. Both assignments were confirmed using our in-house authentic standard library for RT and MS/MS validation (**Fig. 2C**).

We next evaluated MSMICA across 4 independent plasma studies with 3238 samples analyzed under the same LC-HRMS conditions. The benchmark included 160 Schymanski Level 1 metabolites analyzed using hydrophilic interaction chromatography with positive electrospray ionization (HILIC+) and 122 metabolites from reversed-phase chromatography with negative electrospray ionization (C18−). Performance was measured using the fraction correct rate (FCR), defined as the fraction of LC-HRMS features for which the MSMICA identification correctly matched a feature to its associated metabolite (FCR = true positives / all positives). MSMICA achieved an average FCR of 0.994 (0.994 ± 0.004) in HILIC+ and C18− data (**Fig. 2D**). Using the same validation procedure in three serum LC-HRMS datasets, MSMICA achieved an average FCR of 0.990 (0.990 ± 0.005) in HILIC+ and C18− data (**Fig. 2D**).

We further assessed MSMICA by comparing it with other metabolite annotation algorithms, conducting RT validation, performing validation with MS/MS databases, and testing external datasets. For the head-to-head algorithm comparison, we used a human plasma HILIC+ dataset with data-dependent MS/MS acquisition (**Supplementary Data 2**) to compare the performance of different metabolite annotation algorithms using a total of 116 Schymanski Level 1 metabolites served as the benchmark (**Supplementary Data 3**). Using the same feature table and MS/MS input across methods, MSMICA achieved the highest FCR (0.99), the second-highest number of unique metabolite identifications (5655 metabolites), and the highest proportion of confirmed metabolites present in the output (84.5%) (**Fig. 3A**).

**Figure 3.**
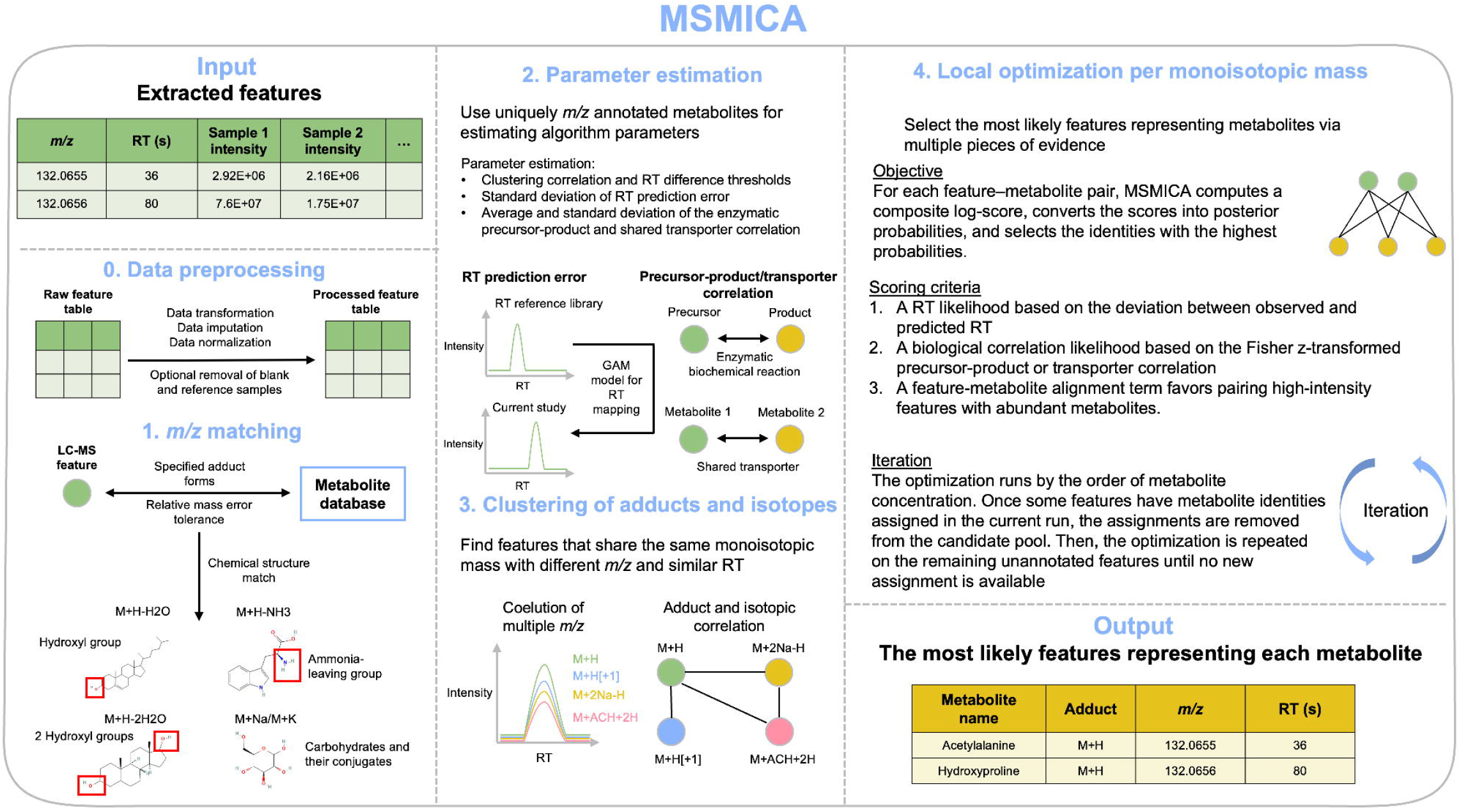
MSMICA additional validation. **(A).** Summary of the fraction correct rates, number of unique metabolites, and percentage of confirmed metabolites among different metabolomics annotation algorithms. The results used a human plasma study HILIC+ data with Schymanski Level 1 identified metabolites (confirmed by 10 ppm *m/z* matching, RT within 30s, MS/MS, and quantification with a check on HMDB for estimated blood concentration when quantification results are ambiguous) as the reference. MS/MS data were used for MetDNA2, MetDNA3, and NetID. xMSannotator used the HMDB metabolite database, while other algorithms used the default metabolite database. For the number of unique metabolites, xMSannotator result was based on HMDB ID, while other algorithms used InChIKey. **(B)**. Summary of the fraction correct rates of MSMICA results in 4 independent plasma LC-HRMS studies and 3 serum LC-HRMS studies using RT identified metabolites (confirmed by 10 ppm *m/z* matching and RT within 30s) as the reference. (**C)**. Summary of the fraction correct rates of MSMICA results in human plasma study with HILIC+, HILIC-, C18+, and C18-data using MS2Query results as the reference. Only the results with MS2Query score ≥ 0.7, theoretical *m/z* within 10 ppm, and RT matched to the feature table within 12 s were used for validation. **(D)**. Summary of the fraction correct rates of MSMICA results in publicly available, external human blood and urine datasets. Metabolites used as the reference were confirmed by the relevant laboratories with the RT and/or MS/MS of authentic standards (Metabolomics Standards Initiative Level 1).

To test performance against a larger RT reference library, we expanded the set to 407 HILIC+ metabolites and 416 C18− metabolites. Across the same 4 plasma datasets and 3 serum datasets, MSMICA achieved average FCR of 0.974, 0.947, 0.988, and 0.991 in plasma HILIC+, plasma C18−, serum HILIC+, and serum C18− datasets, respectively (**Fig. 3B**).

We also evaluated MSMICA using MS2Query^33^ as an external MS/MS matching reference in a study analyzed in four analytical modes: HILIC+, HILIC−, C18+, and C18−. High confidence MS2Query results, defined by a MS2Query score of ≥ 0.7 and theoretical *m/z* within 10 ppm, were used as the reference standard (**Supplementary Data 4**). Under this framework, MSMICA achieved FCR values of 0.933, 0.806, 0.875, and 0.923 in HILIC+, HILIC−, C18+, and C18−, respectively (**Fig. 3C**).

Finally, we tested MSMICA in five external blood and urine datasets generated using LC-HRMS settings different from those used above (**Supplementary Data 5**). In these datasets, MSMICA achieved average FCR values of 0.912 (0.912 ± 0.013) when only Schymanski Level 3a identifications were considered and 0.883 (0.883 ± 0.028) when both Level 3a and Level 3b identifications were included (**Fig. 3D)**.

### MSMICA biological validation using metabolic pathway alterations

In addition to LC-HRMS-based validation, we evaluated MSMICA at the biological pathway level. The rationale is that if a study of a specific disease population has reported pathway-level differences between disease and control groups, then MSMICA should recover many of the same pathway changes in a comparable dataset.

To test this hypothesis, we analyzed a clear cell renal cell carcinoma (ccRCC) dataset containing paired tumor and normal renal tissue samples from seven patients. This dataset was similar to that of a previously published ccRCC study^34^. Metabolites were identified with MSMICA, differential analysis was performed with limma^35^, and KEGG pathway-based analysis was performed with MetaboAnalyst^36^. The analysis identified multiple significant differences between tumor and normal tissue, including changes in glutathione metabolism, amino acid metabolism, and carbohydrate metabolism (**Fig. 4A**).

**Figure 4.**
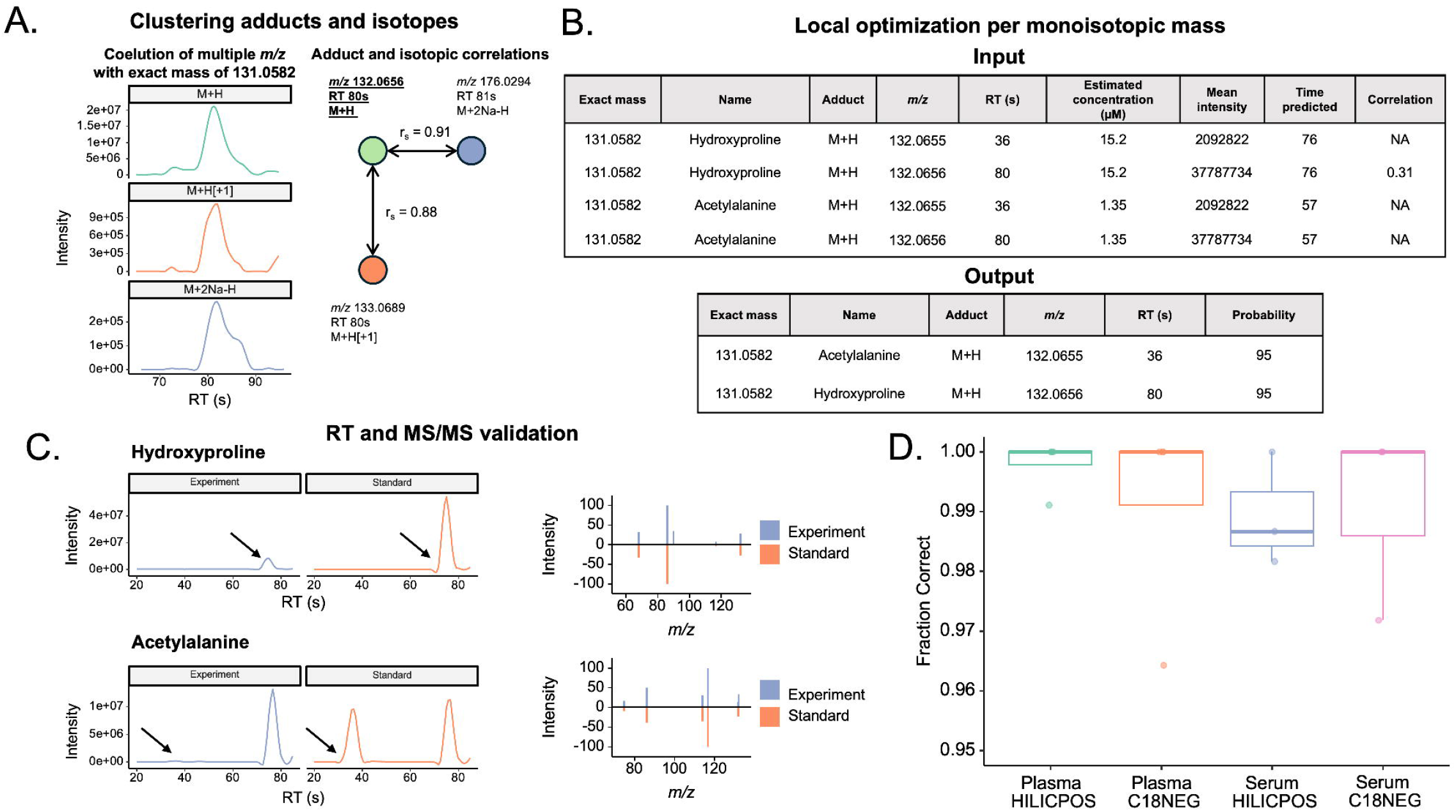
MSMICA biological validation using metabolic pathway alterations. **(A).** KEGG pathway-based analysis of metabolic alterations by comparing ccRCC samples with normal tissue samples. These were paired ccRCC samples and normal kidney tissue samples from 7 patients with ccRCC. Each sample was analyzed by HILIC+ and C18-. Hits ratio is the number of pathway hits divided by the total pathway size. All were found in the previous ccRCC study except folate, linoleic acid, and arginine pathways. The overlapping pathways were bolded. **(B)**. Differential abundance scores reflecting the direction and degree of the metabolic pathway alterations. It is defined as (the number of metabolites increased - the number of metabolites decreased)/the number of all measured metabolites in pathway. A score of 1 demonstrates that all metabolites in the pathway elevate, where -1 demonstrates that all metabolites in the pathway decrease. The overlapping pathways with the same directions are: glycerophospholipid metabolism, purine metabolism, histidine metabolism, alanine, aspartate and glutamate metabolism, glutathione metabolism, beta-alanine metabolism, and pantothenate and CoA biosynthesis. These pathways were also bolded in the figure.

We then compared the pathway results with those from the published ccRCC study^34^. Of the 23 significantly altered pathways identified in our analysis, 20 were also reported previously^34^. We next examined pathway direction using HILIC+ and C18− data. Among pathways with a clear direction of differential abundance, 7 of 10 showed the same direction in both studies^34^ (**Fig. 4B**).

### Coverage of MSMICA results

Because MSMICA can be applied across different LC-HRMS methods and sample types, we tested whether combining results from multiple datasets would increase metabolite identification number and pathway coverage. In a human plasma study, we ran MSMICA separately on HILIC+, HILIC-, C18+, and C18-data and then combined the results by InChIKey. The number of unique identified metabolites increased from 3,886 using HILIC+ alone to 8,500 when all four datasets were combined (**Fig. 5A**).

**Figure 5.**
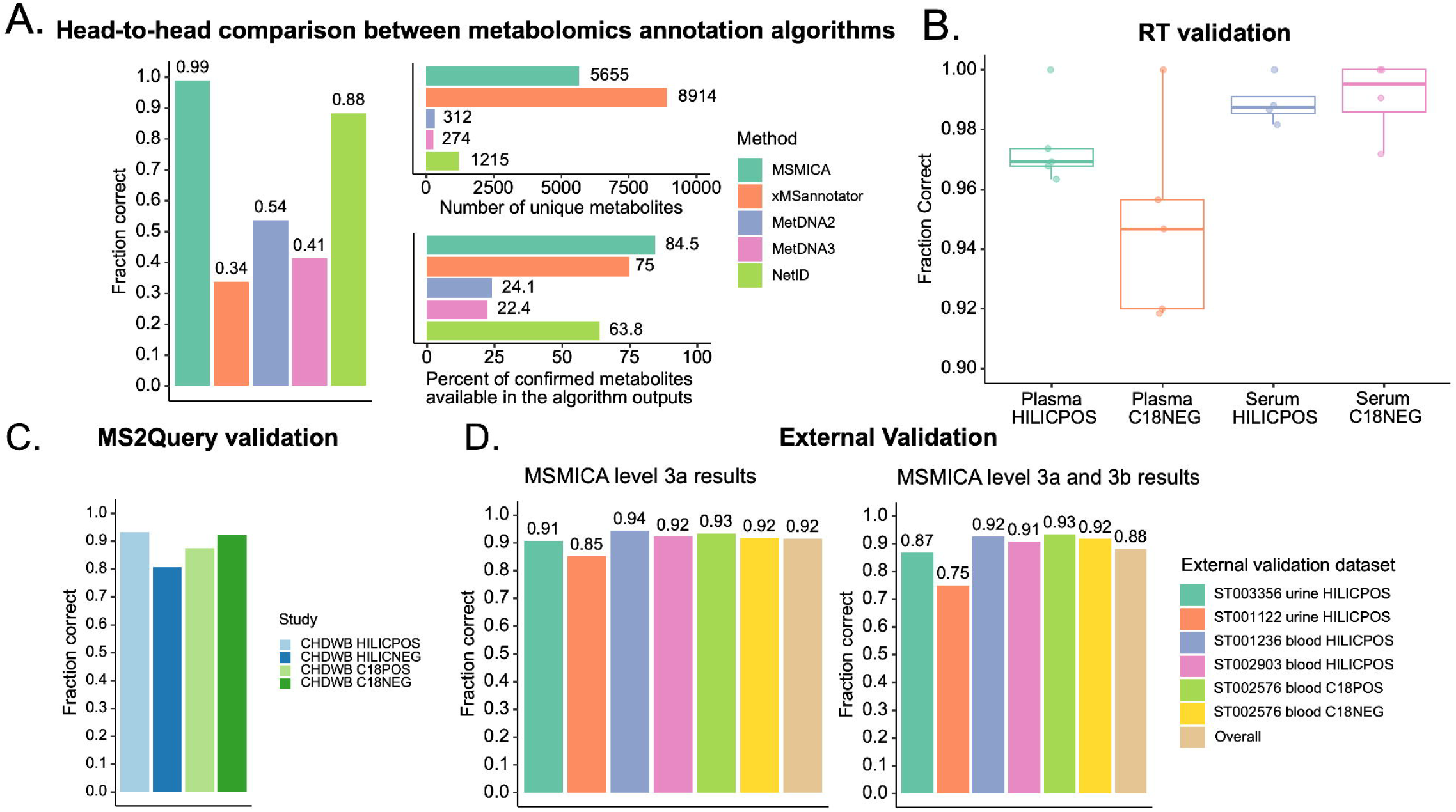
Coverage of MSMICA results for human plasma and mouse tissues. **(A).** The number of unique metabolites increases as the number of analytical methods increases using MSMICA with human plasma LC-HRMS data. The same study samples were analyzed using four different analytical platforms: HILIC+, HILIC-, C18+, and C18-. MSMICA was run for data from each analytical method separately. The number of unique metabolites was calculated using the number of unique InChIKeys (first 14 characters). (**B)**. The number of unique metabolites increases as the number of analytical methods increases using MSMICA with mouse serum, kidney, liver, lung, and testis LC-HRMS data. Data was from an unpublished mouse study. The same mouse samples were analyzed five times using different tissue types: serum, kidney, liver, lung, testis. The number of unique metabolites was calculated using InChIKey. (**C)**. The number of unique metabolites increases as the number of tissue types increases, using MSMICA with LC-HRMS data from five different tissue types with two analytical methods. The same data from Panel **B** were used. (**D)**. The number of metabolites in pathways increases as the number of analytical methods increases. Data from Panel **A** were used. Counts are unique within each method; metabolites detected by more than one method are counted in each corresponding stack segment. The stacked total therefore reflects method-wise detections and is not the number of unique metabolites covering each pathway. (**E)**. The number of metabolites in KEGG pathways increases as the number of analytical methods increases. Data from Panel **B** were used. The counting method is the same as Figure 5D. (**F)**. The number of metabolites in KEGG pathways increases as the number of tissue types increases, using MSMICA with mouse serum and tissue LC-HRMS data. Data from Panel **B** were used. The counting method is the same as Figure 5D.

We performed a similar analysis in a mouse study that included five sample tissue types and two analytical modes from the analysis of the same animals. For each sample type, combining HILIC+ and C18− data increased the number of unique identified metabolites (**Fig. 5B**). When HILIC+ and C18− data were first combined within each tissue type and then combined across tissue types, the number of unique identified metabolites increased from 6,302 in serum alone to 12,742 across all tissue data (**Fig. 5C**).

We then mapped these identified metabolites to KEGG pathways. The coverage gains seen at the metabolite level were also observed at the pathway level, both in the absolute number of metabolites in the pathways (**Fig. 5D, E, F)** and in the percentage of pathway metabolite coverage (**Extended Data Fig. 1**).

### Application of MSMICA for data harmonization

To test whether MSMICA can support harmonization across datasets generated under different LC-HRMS conditions, we applied it to four Alzheimer’s disease (AD) datasets from three published studies: the EFIGA and WHICAP plasma datasets from the AD Knowledge Portal (https://www.synapse.org/Synapse:syn70083418/wiki/635413)^37^, and paired plasma and cerebrospinal fluid datasets from study ST000046 in the Metabolomics Workbench^4^. These datasets include subjects with Alzheimer’s disease and subjects as healthy controls. Metabolite identification and statistical analysis were performed separately for each dataset, and the results were then compared across studies.

This analysis identified 19 metabolites that were consistently decreased in Alzheimer’s disease (**Fig. 6A**) and 20 metabolites that were consistently increased when overlap in at least two studies was required (**Fig. 6B**). In contrast, 22 metabolites showed conflicting directions across studies (**Fig. 6C**). When the significantly altered metabolites were used for KEGG pathway analysis, seven pathways were altered in at least two studies (**Fig. 6D**). Among these, biosynthesis of unsaturated fatty acids, glycerophospholipid metabolism, linoleic acid metabolism, and glycine, serine, and threonine metabolism showed decreases in more than 50% of measured metabolites in the disease group, whereas the other pathways showed mixed patterns.

**Figure 6.**
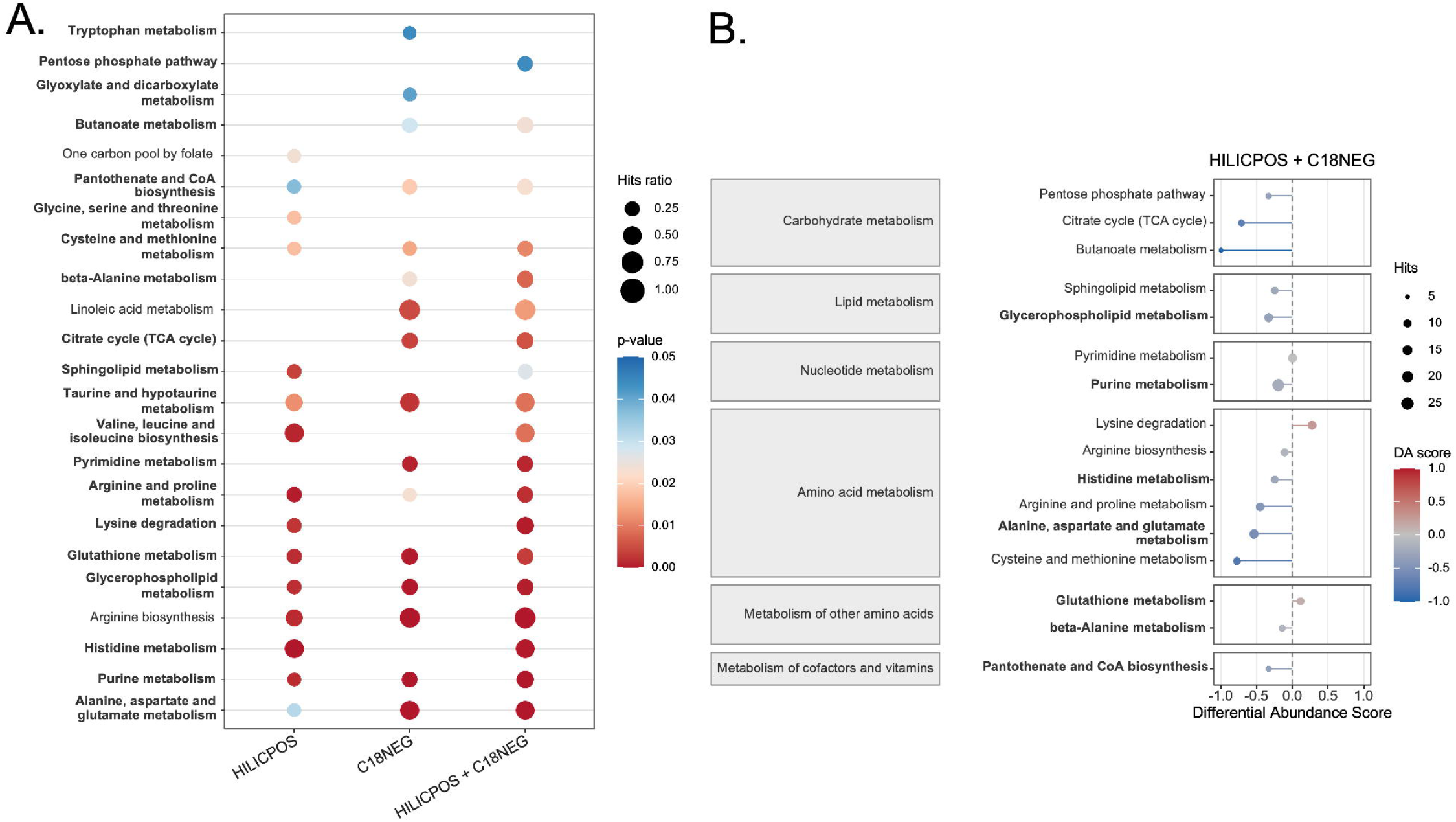
Application of MSMICA for data harmonization in Alzheimer’s disease studies. **(A).** Forest plot of metabolites showing consistent decreases across WHICAP, EFIGA, plasma, and CSF datasets. Only metabolites with a raw p-value < 0.05 were considered. In forest plots, points show log2 fold change and horizontal lines show +/-1.96 standard errors. Log2 fold changes represented the difference between the Alzheimer’s disease group and healthy control group. **(B)**. Forest plot of metabolites showing consistent increases across datasets. **(C)**. Forest plot of metabolites with conflicting directions across datasets. **(D)**. Bubble plot summarizing overlapping pathway-level signals in at least two datasets. In the pathway bubble plot, point size reflects the number of pathway hits and color represents pathway p-value. Duplicated metabolites were removed in the pathway analysis. The Fisher’s Exact Test and the KEGG database were used in MetaboAnalyst’s Pathway Analysis function. HMDB ID of the metabolites were used to maximize the metabolite matches in the analysis. Pathways that appear in at least two studies with a raw p-value < 0.1 were considered.

Sensitivity analysis was performed by adjusting age and sex in the EFIGA and WHICAP studies. The pathway-level findings remained largely consistent, even though some individual metabolite-level results changed (**Extended Data Fig. 2**).

## Discussion

MSMICA integrates mass spectrometry, chromatography, and biological evidence to identify metabolites. After *m/z* matching, MSMICA clusters adduct and isotope features to remove redundant signals originating from the same metabolite. It then applies local optimization within each monoisotopic mass group to iteratively assign metabolite identities to *m/z*-matched features. In this way, MSMICA directly identifies known metabolites without requiring metabolite identities to be assigned to most detected features.

When applied to LC-HRMS untargeted metabolomics data from human biospecimens, MSMICA resolved isomeric metabolites and achieved accurate metabolite identification, as supported by internal and external validation using authentic standard RT and MS/MS data (**Figs. 2 and 3**). At the biological level, MSMICA results for a ccRCC study were comparable to pathway alterations reported in a previous ccRCC study, reproducing 7 of 10 pathways with clear directions of change. Because MSMICA does not require an in-house RT library or study-specific MS/MS data, it can be used broadly across existing LC-HRMS datasets. In our analyses, this flexibility improved metabolite coverage when multiple analytical methods or tissue types were combined (**Fig. 5**) and enabled cross-study comparison and meta-analysis (**Fig. 6**). MSMICA thereby creates opportunities to reanalyze archived data and public metabolomics datasets at both the individual-study and meta-analysis levels.

LC-HRMS untargeted metabolomics typically detects tens of thousands of features, but most are not confidently identified^6^. There is still debate over whether most unannotated features represent novel metabolites or mass spectrometry artifacts^38,39^. Even if only a minority corresponds to known metabolites, confident identification of thousands of known metabolites remains difficult in practice. MSMICA addresses these limitations with offers a new computational approach for metabolite identification. Another common strategy for LC-HRMS metabolite identification is to build metabolic networks in which features are treated as nodes with assigned chemical formulas and edges represent atom transformations between features. NetID advanced this approach for metabolite identification for known metabolites and metabolite discovery for unknown metabolites^17^. However, isomers that share the same formula but differ in RT require MS/MS to differentiate them. In addition, when a feature receives equal NetID scores for multiple candidate identifications, the identification linked to the earliest database ID is reported by default.

MSMICA has several strengths and areas for improvement. Its key strengths are that it does not require RT reference library and MS/MS data inputs and thus offers an automated and flexible approach for computational metabolite identification for many untargeted metabolomics datasets. MSMICA is also optimized for speed with analysis of most metabolomics datasets completed within 10 minutes. Areas for future improvements include the curation of a xenobiotic precursor-product relationship database of drug, food, and environmental toxicant metabolites, as well as support for the identification of unknown metabolites. Thus, the current version of MSMICA cannot be used for metabolite discovery and metabolite identification of certain known xenobiotic metabolites with documented precursor-product relationships. Future research is needed to curate a xenobiotic precursor-product relationship database and support unknown metabolite identification in MSMICA.

Although MSMICA can generate thousands of candidate metabolite identifications, experimental validation of features remains important. Authentic standard RT and MS/MS data provide strong evidence of metabolite identification. Computational methods are particularly useful when they prioritize the most likely metabolite identities for targeted experimental follow-up. MSMICA applies multiple pieces of evidence to substantially improve metabolite identification coverage and accuracy for known metabolites. Overall, MSMICA complements experimental metabolite identification and may help expand the use of metabolomics across a wide range of research applications. MSMICA is an R package available for non-commercial use on GitHub at https://github.com/jamesjiadazhan/MSMICA under the CC BY-NC-ND 4.0 License.

## Methods

### MSMICA parameter estimation

The parameters, including the RT and correlation thresholds for clustering of adducts and isotopes, standard deviation of RT prediction error, and mean and standard deviation of precursor-product/transporter correlation, are obtained during the algorithm run using those annotated metabolites with non-overlapping isomers in chromatography (only one feature in the feature table with a similar *m/z* matched to the metabolites within the *m/z* matching threshold).

The adduct formation, adduct correlation, and isotope correlation probability parameters in the clustering algorithm were estimated using 59 Metabolomics Standards Initiative level 1 metabolites with multiple adduct forms (coelution + 10s of the primary adduct) among 4 independent plasma LC-HRMS studies with 3238 human EDTA plasma samples. The metabolites were identified and quantified by using the RT, MS/MS, and calibration curve of authentic standards^40^.

### Metabolomics dataset description

Four studies with 3238 human EDTA plasma samples, three human and mouse serum datasets, one human plasma dataset with MS/MS data collection in four analytical modes, one human renal tissue dataset, and one mouse tissue dataset, all of which were generated internally, were used. To test algorithm performance with data collected from different metabolomics methods in other laboratories, five external human blood and urine datasets, and three external human plasma and cerebrospinal fluid (CSF) datasets related to Alzheimer’s disease were used in this manuscript. All the studies were approved by the Institutional Review Board (IRB) of Emory University or other collaborators’ institutions.

#### Internal LC-HRMS datasets

- The four human plasma studies were analyzed by the same LC-HRMS setting across five years^41– 44^.
- Serum studies consisted of a human dataset containing 650 serum samples from African American pregnant women^45^, 1148 serum samples from pregnant women^46^ (Metabolomics Workbench ST005056), and 40 serum samples from a longitudinal observational study in mice^47^.
- The human plasma dataset from generally healthy subjects with MS/MS data collection had a sample size of 715 for C18+ and C18- and a sample size of 357 for HILIC+ and HILIC-^48^.
- The human renal tissue dataset includes 7 renal tumor samples paired with 7 adjacent normal renal tissue samples from 7 patients with renal cell carcinoma (Emory IRB protocol 2025P012499).
- The tissue study included mice with five different tissue types collected (n=48).
- Within each study, in-house pooled reference human plasma and US National Institute of Standards and Technology (NIST) SRM-1950 human pooled plasma samples were used for quality control and metabolite reference quantification.

#### External LC-HRMS datasets

- Five external blood and urine validation datasets were downloaded from the Metabolomics Workbench (https://www.metabolomicsworkbench.org/ with study IDs ST003356, ST001122, ST001236, ST002903, and ST002576.
- For the datasets related to Alzheimer’s disease, EFIGA and WHICAP plasma datasets were downloaded from https://www.synapse.org/Synapse:syn70083418/wiki/635413, and the other plasma and CSF datasets were downloaded from the Metabolomics Workbench with the study ID of ST000046.

### Metabolomics sample preparation

For studies in which data are first reported here, fluid samples were mixed with an ice-cold acetonitrile-internal standard solution at a 1:2 (sample: solution) ratio, incubated on ice, centrifuged, and the supernatants were collected for LC-MS analysis, using protocols similar to those previously described^40^. Mouse serum from one study^47^ was extracted using a solution containing added water to account for low serum volumes.

For renal cell carcinoma samples (and paired normal kidney samples), 20□μl of an ice-cold mixture containing 2:1 acetonitrile:water with an internal standard was added per milligram of tissue, and samples were then minced with iris scissors. Samples were then subjected to additional homogenization using a Dounce glass pestle. Tissue samples were then vortexed, kept on ice, and placed on an orbital shaker for 30 minutes. Following this, samples were briefly vortexed again, then centrifuged at 14,000 rpm for 20 minutes at 4 °C to precipitate proteins. The supernatants were then collected, transferred into autosampler vials, and stored at −70□°C until analysis. For all the other tissue samples, 15□μl of an ice-cold mixture containing 2:1 acetonitrile:water with an internal standard was added per milligram of tissue, followed by homogenization using a handheld pellet pestle. The samples were then vortexed and kept on ice for 30 minutes before undergoing centrifugation at 14,000□g for 10 minutes at 4□°C to precipitate proteins. The supernatants were collected into autosampler vials and stored at −70□°C until they were analyzed instrumentally. For data obtained from public access databases, sample processing descriptions can be found together with the data.

### Data processing

LC-MS raw files were converted to mzXML profile format and extracted to create feature tables (with *m/z*, RTs, and intensities) using apLCMS^49^ and xMSanalyzer^50^. For public access datasets, if feature tables were directly available, they were used by the MSMICA algorithm to generate outputs. If not, the LC-MS raw files were converted to mzML centroid format and extracted to build feature tables using asari^51^.

To prepare MS/MS data for MS2Query and METDNA2, MZmine (version 4.0.8) was used for feature extraction and to pair MS/MS scans with MS^1^ scans and generate .mgf files^52,53^. MS2Query results were filtered with thresholds of MS2Query score ≥ 0.7 and the *m/z* difference between sample precursor *m/z* and library precursor *m/z* ≤ 10 ppm. MS2Query results were then filtered by matching to the feature table with thresholds of *m/z* ≤ 10 ppm and RT 12 s. The mgf file generated by MZmine in HILIC+ was used in the METDNA2, METDNA3, and NetID algorithms.

### MSMICA setting

A mass error threshold of 10 ppm is used for *m/z* matching in all studies. With high-resolution instruments and adequate calibration, a much lower threshold (3-5 ppm) can be used. However, to ensure interoperability of MSMICA testing across the several datasets used, with varying degrees of mass accuracy, a less stringent 10 ppm threshold was used. Positive adduct lists for the MSMICA clustering algorithm were “M+H”, “M+Na”, “M+2Na-H”, “M+H-H2O”, “M+H-NH3”, “M+ACN+H”, “M+ACN+2H”, “2M+H”, “M+2H”, “M+H-2H2O” for plasma, serum, and urine samples and “M+H”, “M+K”, “M+2K-H”, “M+H-H2O”, “M+H-NH3”, “M+ACN+H”, “M+ACN+2H”, “M+ACN+K”, “2M+H”, “M+2H”, “M+H-2H2O” for tissue samples. Negative adduct lists for the MSMICA clustering algorithm were “M-H”, “M+Cl”, “M+FA-H” (formic acid adduct), “M+Hac-H” (acetic acid adduct), “M+FA+Na-2H”, “M+Na-2H”, “M-2H”, “2M-H”, “M+ACN-H” for plasma, serum, and urine samples and “M-H”, “M+Cl”, “M+FA-H”, “M+Hac-H”, “M+FA+K-2H”, “M+K-2H”, “M-2H”, “2M-H”, “M+ACN-H” for tissue samples.

### MSMICA chemical structure rules

Two families of rules constrain which adducts are allowed for each metabolite, plus one post□hoc exclusion to avoid *m/z* coincidence between structurally related compounds.

1. Functional□group rules

a. M+H-NH3 adduct is only considered when the metabolite has at least one free amine group.
b. M+H-H2O adduct is only considered when the metabolite has at least one carboxyl or hydroxyl group.
c. M+H-2H2O adduct, it is only considered when the metabolite has at least two carboxyl or hydroxyl groups.

2. Chemical subclass□based rules

a. All metabolites prefer M+H or M-H unless the following chemical subclass are observed.

The chemical subclass is computed by ClassyFire.

i. Carbohydrate and carbohydrate conjugates prefer M+Na, M+K, and M+Cl
ii. Bile acids, alcohols and derivatives prefer M+H-H2O and M+H-2H2O
iii. Retinoids prefer M+H-H2O
iv. Pregnane steroids prefer M+H-H2O and M+H-2H2O
v. Gluconic-acid-like compounds prefer M+Na, M+K, M+2Na-H, M+2K-H

From the clustering results, secondary adducts and isotopes were excluded from later analysis to avoid their confusion with the primary adduct of other metabolites. However, even some metabolites have M+H-H2O, M+H-2H2O, and M+H-NH3 adducts in the clustering results, these features are not removed from later analysis because these adducts frequently coincide with M+H of chemically similar metabolites (e.g., 3□hydroxydecanoyl carnitine M-H2O+H vs. decenoylcarnitine C10:1 M+H).

### Data analysis

For Figures 4 and 6, MSMICA results were analyzed by using the limma algorithm^35^. Missing intensity data were imputed using the half-minimum method^54^. Log_2_ fold changes were calculated by comparing the control and disease groups.

### Differential Abundance Score

Differential abundance scores reflecting the direction and degree of the metabolic pathway alterations. After determining which metabolites are significantly altered between the disease and control groups, the differential abundance score is defined as

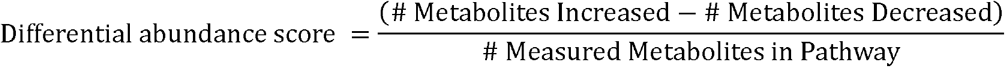

### Reference library

The authentic chemical standards used to build our reference library were from the Sigma-Aldrich Mass Spectrometry Metabolite Library (manufactured by IROA technologies) with stated purities of > 95%. Features associated with potentially formed adducts (M+H, M+2Na-H, M+Na, M-H2O+H, M+K, M+3H, M-2H2O+H, 2M+H, 2M-H, and 2M+ACN+H in HILIC+; M-H, M+Cl, M+Hac-H, M+FA-H, M-2H, 2M-3H in C18-) were checked for each metabolite using a 3 ppm mass window in Thermo xCalibur (v.4.2) QualBrowser software for feature area increases and expected MS/MS patterns after adding standards^40^.

### Metabolite database

The default metabolite database was created by merging the HMDB database^32^ with the KEGG database^31^ manually using InChIKey, SMILES, and metabolite name (**Supplementary Data 1**). Only the metabolites with dominant chiral structure were used; L-sugars, D-amino acids, and a few other metabolites (e.g., D-Carnitine) were removed using ClassyFire^55^.

### Retention time reference library database

The RT reference library used in the MSMICA algorithm for RT mapping and prediction was built by merging the metabolites with experimentally measured RT from the HILIC plasma dataset provided by Retip^56^ (970 metabolites in 17-min LC-MS) and the C18 RIKEN dataset (494 metabolites in 15-min LC-MS), and the metabolites in the default metabolite database with predicted RT using Retip^56^ and RT Pred^57^. Where experimentally measured RT was available, predicted RT was not used. This database is available as **Supplementary Data 6** and can be found in the MSMICA package as hmdb_metabolites_reference_retention_time.rda.

### Metabolite concentration reference database

The biospecimen-specific metabolite concentration reference database was mainly generated using the HMDB database and a recent paper reporting quantification data from the NIST SRM 1950 human pooled plasma^58^. When multiple metabolite concentration values were available from different sources, their averages were calculated. Only metabolite concentration values from adults were included to avoid abnormal values from other populations, such as those with inborn errors of metabolism. When metabolites were not quantified in the literature but metabolite concentrations in healthy adult populations were available from laboratory records, we appended these metabolite concentration values to the database. This database is available as **Supplementary Data 7** and can be found in the MSMICA package as hmdb_metabolites_concentrations_average.rda.

### Metabolic precursor-product relationship database

The biochemical precursor-product reaction database was created by merging the KEGG^31^, BKMS-react^59^, rhea^60^, and Recon3D^61^ databases. In the KEGG database, only the reactions catalyzed by human enzymes were kept. In the Recon3D database, only the human and gut microbiome reactions were kept. This database is available as **Supplementary Data 8** and can be found in the MSMICA package as Recon3D_BKMS_react_rhea_KEGG_connection_mammalia.rda.

### Metabolite co-transporter relationship database

The metabolite co-transporter relationship database was created by merging the Recon3D^61^, UniProtKB^62^, and the database created by Deo et al.^63^ from a literature search. This database is available as **Supplementary Data 9** and can be found in the MSMICA package as Recon3D_unitprot_Deo_human_transporter.rda.

### MSMICA Workflow

The workflow for each step of MSMICA is detailed as follows:

#### Step 0-data preprocessing

Optional sample metadata are used to retain only study samples by removing the reference and blank samples. Feature intensities are log2-transformed, missing values are imputed, and sample-wise quantile normalization is applied. Missing data imputation was performed for features that appeared in less than 20% of all samples using the quantile regression imputation of left-censored data (QRILC) approach^54^ for feature tables with triplicates of each sample and the half minimum method for feature tables with single injection for each sample. Duplicate features with the same rounded *m/z* (4 decimal places) and RT (integers) are collapsed by keeping the signal with the highest mean intensity. In parallel, within the metabolite database, duplicate structures were consolidated at the level on the first 14 characters of the InChIKey.

#### Step 1-*m/z* matching

The workflow begins with a feature intensity table as the primary input. User-provided adduct forms are used to calculate the *m/z* for each metabolite with different adduct forms based on the monoisotopic mass in the metabolite database (**Supplementary Data 1**). The feature table’s *m/z* is matched to the database *m/z* using a specified relative mass error (default 10 ppm), returning all *m/z* matched results. Chemically implausible adducts are removed using chemical structural rules, such as requiring water-loss adducts only for compounds with hydroxyl or carboxyl groups and ammonia-loss adducts for compounds containing an ammonia-leaving group. This yields an initial candidate list for both regular adducts and major isotopologue features.

#### Step 2-Algorithm parameter estimation

MSMICA uses annotated metabolites with a protonated/deprotonated adduct form, non-overlapping isomers in chromatography (only one feature in the feature table with a similar *m/z* matched to the metabolites within the *m/z* matching threshold) to estimate parameters used in the assessment of clustering of adducts and isotopes, RT prediction, and precursor-product/transporter relationship. We hereafter refer to these annotated metabolites as “anchor” metabolites. RTs of all the metabolites in the metabolite database are either experimentally measured or predicted by Retip^56^ and RT Pred^57^ using experimental data as the training dataset (HILIC: 970 metabolites from the MassBank of North America database, C18: 494 metabolites from the RIKEN plant specialized metabolome annotation database). The predicted RTs of metabolites in the reference RT library were first mapped to the “anchor” annotated metabolite using a monotonically constrained generalized additive model (GAM) adapted from PredRet.^64^ The fitted GAM model is then used to predict expected RTs for all candidate metabolites in the current study, and the residual standard deviation provides an empirical RT uncertainty parameter for downstream scoring. Then, the same “anchor” metabolites are used to estimate the average and standard deviation of precursor-product/transporter correlations by conducting correlations among themselves.

Similarly, the same metabolites with protonated/deprotonated adduct forms are used to estimate the RT difference and correlation thresholds among the same metabolites with multiple adducts. Based on our investigation of adduct and isotopic correlation patterns in the 4 in-house plasma LC-HRMS studies, only adducts and isotopes with a minimum correlation of 0.4 and the adducts with a RT difference less than 15 s and isotopes with a RT difference less than 10 s were kept. Then, for the study where MSMICA is used, RT and Spearman correlation coefficient thresholds are estimated based on the thresholds that included 80% of adducts from the same metabolites in the study.

#### Step 3-Clustering adducts and isotopes

This step resolves cases in which multiple features share the same monoisotopic mass. It selects the most likely features representing metabolites with the same monoisotopic mass via clustering of adducts and most abundant isotopologues. Features supported by adduct correlation and/or isotopic correlation are then scored with a Bayesian model that combines three binary evidence types: whether the feature is a common primary adduct, whether it belongs to an adduct-correlation cluster, and whether it has a correlated isotopologue. Posterior probabilities are normalized within each monoisotopic-mass cluster, and the highest-probability features are retained as the most likely representatives of that metabolite signal cluster. This step is used to identify robust features of metabolites and to exclude correlated secondary adducts from later rounds of identification.

##### 1. Cluster definition

For a given monoisotopic-mass group, let the candidate feature set be

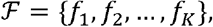

where *f*_*i*_ denotes a matched LC-MS feature assigned to the same monoisotopic mass as others in the set, *F*. The objective is to estimate which feature is the most likely representative ion for that cluster.

##### 2. Prior probability

Before considering evidence, MSMICA assigns a uniform prior over the *K* candidate features:

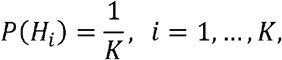

where *H*_*i*_ denotes the hypothesis that feature *f*_*i*_ is the true representative feature in the cluster.

##### 3. Evidence variables

For each feature *f*_*i*_, three binary evidence variables are defined.

###### 3.1 Adduct type

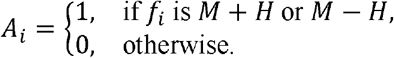

###### 3.2 Adduct correlation

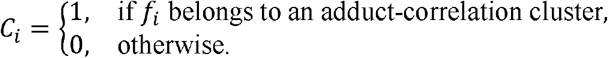

A correlated feature pair (*f*_*i*_, *f*_*k*_)is declared when

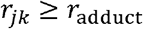

where *r*_*adduct*_ is the adduct correlation coefficient threshold, and *r*_*jk*_ is the Spearman correlation between feature j and feature k.

and

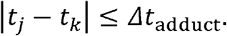

where Δ*t*_adduct_ is the retention time difference between feature j and feature k.

###### 3.3 Isotope-support correlation

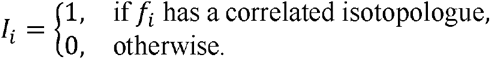

For a primary feature *f*_*i*_ and its candidate isotopic partner 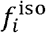 with the same monoisotopic mass, the sample-wise abundance ratio is

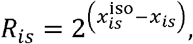

where *x*_*is*_ and 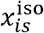 are the log2-transformed intensities in sample *s*.

The mean abundance ratio is

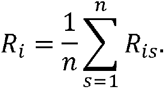

An isotopic pair is accepted when the mean 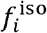 abundance is not higher the mean *f*_*i*_, and their Spearman correlation passes the isotopic correlation threshold

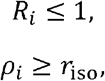

where *r*_iso_ is the isotopic correlation coefficient threshold and *ρ*_*i*_ is the Spearman correlation between the primary feature *f*_*i*_ and its candidate isotopic partner 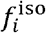

and

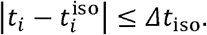

where Δ*t*_iso_ is the retention time difference between *t*_*i*_and 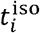.

##### 4. Empirical conditional probabilities

The algorithm estimates evidence probabilities under two conditions:

- *H*: the feature is the correct representative feature of the metabolite that is validated by RT and MS/MS of authentic standards
- ⌝*H*: the feature is not the correct representative feature of the metabolite that is validated by RT and MS/MS of authentic standards

As indicated above, the following adduct formation, adduct correlation, and isotope correlation parameters in the clustering algorithm were estimated using 59 Metabolomics Standards Initiative level 1 metabolites with multiple adduct forms (coelution + 10s) among 4 independent plasma LC-HRMS studies with 3238 EDTA human plasma samples.

For adduct type evidence,

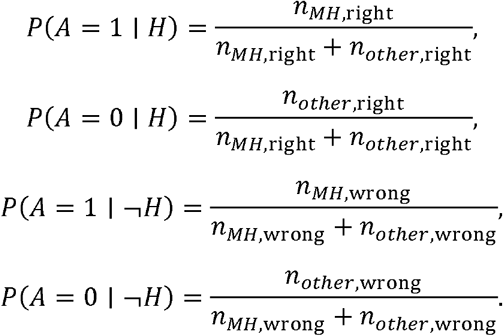

Where *n*_*MH*,right_ estimates how likely features are correct when protonated or deprotonated adduct is present (default 112/147), *n*_*MH*,wrong_ estimates how likely features are wrong when protonated or deprotonated adduct is present (default 35/147), *n*_*other*,right_ estimates how likely features are correct when the other forms of adducts are present (default 97/476), and *n*_*other*,wrong_ estimates how likely features are wrong when the other forms of adducts are present (default 379/476).

For adduct-correlation evidence,

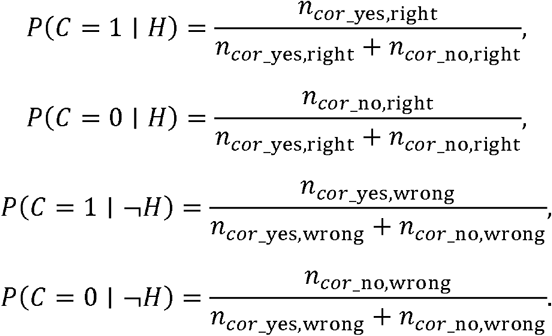

Where *n*_*cor*_yes,right_ estimates how likely features are correct when adduct correlation is present as defined by Spearman correlation coefficient ≥ 0.39 and retention time difference ≤ 6 s (default 90/158), *n*_*cor*_yes,wrong_ estimates how likely features are wrong when adduct correlation is present (default 68/158), *n*_*cor*_no,right_ estimates how likely features are correct when adduct correlation is not present (default 119/465), and *n*_*cor*_no,wrong_ estimates how likely features are wrong when the adduct correlation is not present (default 346/465). The Spearman correlation coefficient and retention time difference thresholds were estimated by using the thresholds that capture 80% of features with experimentally validated adduct correlations in the 4 plasma LC-HRMS studies with 3238 plasma samples.

For isotope evidence,

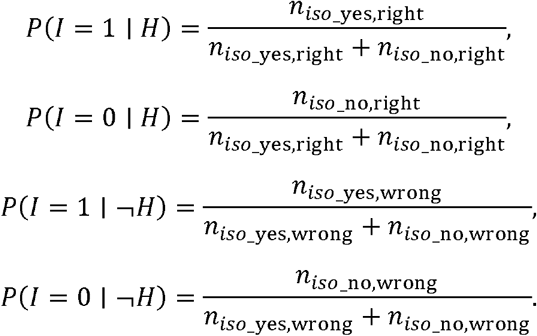

Where *n*_*iso*_yes,right_ estimates how likely features are correct when isotopic correlation is present as defined by Spearman correlation coefficient ≥ 0.71 and retention time difference ≤ 4 s (default 43/61), *n*_*iso*_yes,wrong_ estimates how likely features are wrong when isotopic correlation is present (default 18/61), *n*_*iso*_no,right_ estimates how likely features are correct when isotopic correlation is not present (default 212/876), and *n*_*iso*_no,wrong_ estimates how likely features are wrong when the isotopic correlation is not present (default 664/876). The Spearman correlation coefficient and retention time difference thresholds were estimated by using the thresholds that capture 80% of features with experimentally validated adduct correlations in the 4 plasma LC-HRMS studies with 3238 plasma samples.

##### 5. Log-likelihood ratios for each evidence type

For each feature *f*_*i*_, each evidence source is converted into a log-likelihood ratio.

###### 5.1 Adduct evidence

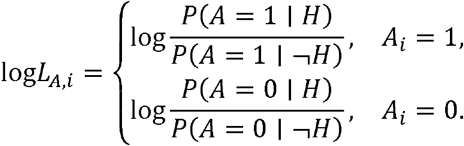

###### 5.2 Adduct-correlation evidence

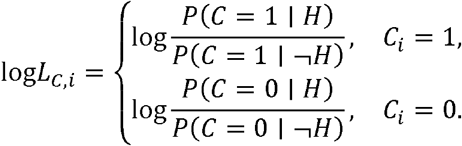

###### 5.3 Isotope evidence

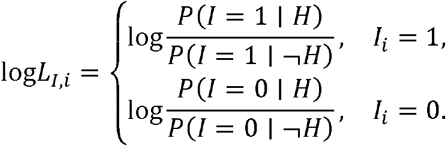

##### 6. Posterior scoring in log space

For each candidate feature *f*_*i*_, MSMICA forms an unnormalized log-posterior score:

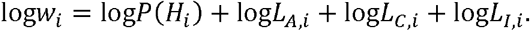

With the uniform prior,

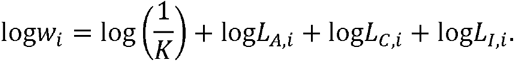

##### 7. Posterior probability normalization

Posterior probabilities are obtained by softmax normalization within the cluster:

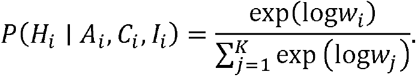

For numerical stability, the implementation uses centered scores:

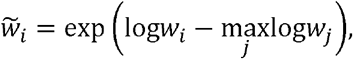

followed by

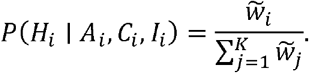

The reported percentage is

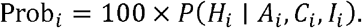

##### 8. Final identification rule

A feature is selected by the clustering algorithm if it attains the maximum posterior probability within the cluster:

##### 9. Special case

If only one candidate feature remains after filtering, it is assigned directly:

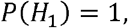

which is equivalent to

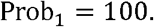

##### 10. Summary

The core clustering model can be written as

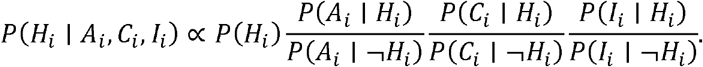

Equivalently, in log form,

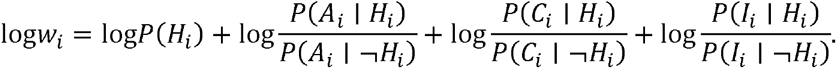

Normalization across the *K* candidate features gives

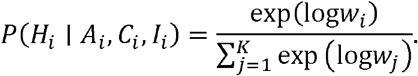

##### 11. Assumptions

Biospecimens contain well-characterized metabolites with known monoisotopic mass, highly abundant metabolites are present in LC-HRMS with multiple isotopologues and adduct forms, and adduct correlation, isotopologue correlation, and adduct formation are assumed to have minimal dependency on each other. Therefore, computational approaches can be used to obtain accurate mass and likely identifications for highly abundant metabolomics features. Based on the assumptions, the algorithm hypothesizes that features with adduct correlation, isotope correlation, and preferred adduct formation are more likely to represent the corresponding metabolites than other features.

#### Step 4-Local optimization per monoisotopic mass

After *m/z* matching, RT mapping, precursor-product or shared-transporter correlation scoring, and feature intensity metabolite concentration alignment, MSMICA applies local optimization within each monoisotopic-mass group. This step resolves cases in which multiple LC-MS features and multiple candidate metabolites share the same monoisotopic mass. The current implementation uses a metabolite-first greedy assignment strategy: Metabolites are evaluated from highest to lowest expected biospecimen concentration, and each metabolite selects at most one still-available LC-MS feature from the same monoisotopic-mass group.

##### 1. Feature-metabolite pairs

For each monoisotopic-mass group, MSMICA resolves ambiguous feature-metabolite assignments by evaluating all candidate feature-metabolite pairs within the group and integrating orthogonal evidence in a scoring framework. Let

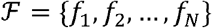

be the set (N) of observed LC-MS features assigned to a monoisotopic mass, and let

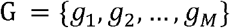

be the set (M) of candidate metabolites associated with that same mass group.

Each possible assignment is represented by a feature-metabolite pair

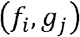

where *f*_*i*_ is an observed feature and *g*_*i*_ is a candidate metabolite. Therefore, each mass group contains N * M possible assignments. If there are 2 features matched to 2 metabolites with the same mass based on *m/z*, there are 4 possible assignments. For example, for feature 1 *m/z* 132.0655 RT 36 s and feature 2 *m/z* 132.0656 RT 80 s, if only hydroxyproline and acetylalanine are considered as candidate metabolites, then there are 4 possible assignments: feature 1 annotated as hydroxyproline, feature 2 annotated as hydroxyproline, feature 1 annotated as acetylalanine, and feature 2 annotated as acetylalanine.

For each candidate pair (*f*_*i*_, *g*_*i*_), the algorithm computes a composite log-score based on RT prediction agreement, precursor-product or shared-transporter correlation, and a feature intensity metabolite concentration alignment. The feature intensity metabolite concentration alignment favors the feature-metabolite pair in which relatively high intensity features are paired with relatively abundant metabolites in the algorithm’s specified biospecimen (default is blood).

##### 2. Feature and metabolite notation

For the pair (*f*_*i*_, *g*_*j*_), let ΔRT_*ij*_ be the absolute retention-time difference between the observed RT of the feature and the predicted RT of the metabolite,

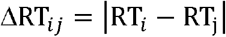

where RT_*i*_ is the observed RT of feature *f*_*i*_ and RT_*j*_ is the mapped RT predicted for metabolite *g*_*j*_.

If a biological correlation is available for the pair, let *r*_*ij*_ denote the Spearman correlation coefficient between the sample intensity of feature-metabolite pair *f*_*i*_*g*_*i*_ and the sample intensity of its connected precursor, product, and transporter-related metabolite.

The empirical mean and standard deviation of the correlation coefficients among high-confidence training pairs are denoted by *μ*_corr_ and *σ*_corr_, respectively. The standard deviation of the retention-time residuals from the study-specific RT mapping model is denoted by *σ*_RT_.

##### 3. RT and correlation modeling

MSMICA supports two calibration modes, “current” and “empirical” (default), for converting retention-time and correlation evidence into log-likelihood or log evidence score contributions. In the “current” parametric mode, retention-time evidence is modeled with a Gaussian error model,

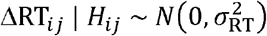

where the mean RT difference is 0 second, and the RT difference variance is 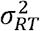. After log transformation, the corresponding log-likelihood is

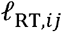

When valid precursor-product or shared-transporter correlation evidence is available, the correlation evidence is modelled as

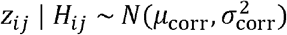

where mean is *μ*_corr_ mean of the Fisher-transformed Spearman correlation coefficient, and variance is 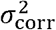. Precursor-product and shared-transporter correlations are all considered..

After log transformation, the corresponding log-likelihood is

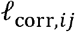

If no precursor-product or shared-transporter correlation is available for the pair, MSMICA assigns a neutral contribution.

In the “empirical” calibration mode (default), MSMICA first evaluates calibrated empirical functions for retention-time residuals and correlation values:

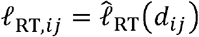

and

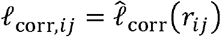

These calibrated functions are estimated from the evidence-calibration object generated upstream while estimating the algorithm parameters. These quantities represent empirical percentile-based support scores rather than probability densities. Missing correlation evidence remains neutral.

Let *ℓ*_RT,*ij*_ and *ℓ*_corr,*ij*_ denote the effective retention-time and correlation log-likelihood contributions after applying the selected calibration mode and any necessary fallback. The pairwise log-score without considering metabolite concentration evidence is then

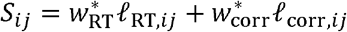

where *w*_RT_ and *w*_corr_ are user-defined evidence weights, and *ℓ*_RT,*ij*_ and *ℓ*_corr,*ij*_ are the likelihood functions of the retention time and correlation evidence. *w*_RT_ and *w*_corr_ default values are 1.

##### 4. Feature intensity metabolite concentration alignment

For each metabolite *g*_*j*_, its expected biospecimen-specific concentration is obtained from HMDB. To prevent all features with the same mass from being assigned to the most abundant metabolites if one metabolite concentration is much higher than the other metabolite concentrations, we use feature intensity metabolite concentration alignment instead of metabolite concentration.

The feature intensity metabolite concentration alignment is based on observed feature intensity and expected metabolite concentration. This term favors assignments in which relatively high intensity features are paired with relatively abundant metabolites.

Within each mass group, features are ordered by decreasing observed intensity and metabolites are represented by their expected concentration. Let **u**_*i*_ be the min-max scaled mean intensity of feature *f*_*i*_, and let **v**_*j*_ be the corresponding min-max scaled estimated metabolite concentration of metabolite *g*_*j*_.

The min-max scale is completed as

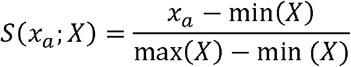

The alignment for candidate pair (*f*_*i*_, *g*_*i*_) is then defined as

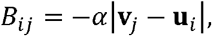

where *α* controls the strength of the alignment. The default α is 1. The alignment prioritizes stronger features to be paired with more abundant metabolites continuously.

The adjusted score becomes

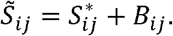

Which can be also described as

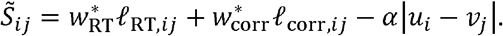

##### 5. Metabolite-first greedy assignment

Within each monoisotopic-mass group, MSMICA assigns features to candidate metabolites using a local optimization greedy approach. Candidate metabolites are processed in decreasing order of their expected biospecimen-specific concentration. Then, it prioritizes the pairing of features with higher intensities with abundant metabolites. This is similar to NetID that aims to find the condition where the global node and edge scores are maximized^17^ while the MSMICA local optimization greedy approach aims to find the feature with the maximized probability of being a given metabolite among the features matched to the same mass.

Let *A*_*j*_ denote the set of available candidate features for metabolite *g*_*j*_. For each available feature *f*_*i*_ ∈ *A*_*j*_, the adjusted score 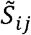 is converted to a normalized assignment probability:

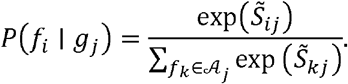

The selected feature is the available feature with the highest assignment probability:

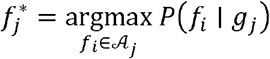

After a feature 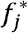 is selected for *g*_*j*_, that feature is removed from the available feature set. The algorithm then evaluates the next metabolite. This procedure enforces one-to-one matching within each local mass group, so each metabolite can select at most one feature, and each selected feature can be used only once.

##### 6. Iterative optimization

After local optimization is completed for all monoisotopic-mass groups for the most abundant metabolites in the current iteration, the retained assignments from the current iteration are stored. MSMICA then removes from the remaining candidate pool any row whose feature coordinate or metabolite InChIKey has already been accepted. The next iteration starts for all monoisotopic mass groups for the second most abundant metabolites. The iterative procedure stops when no input candidates remain or when no new local optimization result is found.

Because some metabolites with very close but non-identical monoisotopic masses can match the same rounded LC-MS feature coordinate (e.g. butyrobetaine and acetylcholine), MSMICA applies an additional cross-mass concentration tiebreaking step. For each observed feature coordinate, represented by *m/z* and RT, if multiple selected metabolites remain, the candidate or candidates with the highest estimated biospecimen-specific concentration are retained.

##### 8. Summary

Overall, MSMICA uses a greedy assignment approach that scores each feasible feature-metabolite pair using retention-time, correlation, and rank-alignment evidence for features matched to the same mass, then sequentially assigns each metabolite to the highest-scoring available feature while enforcing one-to-one feature-metabolite matching. The approach proceeds in the order of metabolite concentration. Once features are identified as metabolites in one iteration, they are removed from the candidate pool, and the next iteration starts until no new metabolite identifications are available.

##### 9. Assumptions

First, because metabolic pathways are regulated at specific sites, metabolites have biochemical reactions catalyzed by enzymes, and many metabolites share the same transporters, the intensities of metabolites that are precursors or products of each other or share the same transporters tend to be correlated. Therefore, precursor-product/transporter correlation can be used to support the metabolite identity of features. Second, the algorithm assumes that even for the same products, each precursor-product reaction pair has a limited dependency on the other precursor-product reaction pair. Based on the assumptions, the algorithm hypothesizes that if the features are indeed the correct metabolites, they should have a higher correlation coefficient than the other features. Third, the algorithm assumes that the average biospecimen-specific concentration from HMDB can be used for building biospecimen-informed concentration contribution because literature-supported abundant metabolites in the same biospecimen type are expected to be consistently abundant in the current study if the same type of biospecimen is used and there is no extreme abnormal metabolism.

### Final Output

The final output is a table of metabolites with database identifiers, structural descriptors, adduct assignment, observed and predicted RT, posterior probability, feature intensity, and the evidence types supporting each identification. Evidence is tracked in the output as a structured identification method string, indicating whether a metabolite was supported by *m/z* matching alone or additionally by RT prediction, biospecimen-specific concentration, precursor-product/transporter correlation, and clustering of adducts and isotopes. An optional companion table includes secondary adduct and isotopic features associated with the identified metabolites. Features not identified by the workflow, including those with no *m/z* matches to the HMDB and KEGG database, can also be exported for downstream manual inspection or future analysis.

### Liquid chromatography and high resolution mass spectrometry

Extracts were analyzed using liquid chromatography (Thermo Scientific Dionex Ultimate 3000) coupled with Orbitrap high-resolution mass spectrometry (Thermo Scientific HF-Q Exactive or ThermoFisher Orbitrap Fusion Tribrid). The chromatographic system has a dual pump configuration for parallel analyte separation and column washing^40^. Sample extracts were analyzed by HILIC+ and C18-. HILIC consisted of a Waters XBridge BEH Amide XP HILIC column (2.1 × 50 mm^2^, 2.6 μm particle size) and mobile phases A (LCMS-grade water), B (LCMS-grade acetonitrile), and C (2% formic acid). The gradient started with 22.5% A, 75% B, and 2.5% C for the first 1.5 minutes, increased linearly to 75% A, 22.5% B, and 2.5% C to 4 minutes, and then held for 1 minute. C18 chromatography consisted of a Higgins Targa C18 column (2.1 × 50 mm^2^, 3 μm particle size) and phases A (water), B (acetonitrile), and C (10 mM ammonium acetate). The gradient started with 60% A, 35% B, and 5% C for the first minute, increased linearly to 0% A, 95% B, and 5% C for 3 minutes, and then held for 2 minutes. For both methods, the flow rate was set to 0.35 mL/min for the first minute, then increased to 0.4 mL/min for the final 4 minutes.

For the CHDWB human plasma study used for the head-to-head comparison between metabolomics annotation algorithms and MS2Query validation, samples were analyzed using a Thermo Scientific Vanquish Duo HPLC coupled to a Thermo Scientific ID-X mass spectrometer in HILIC+, HILIC-, C18+, C18-. The HILIC method consisted of a Waters ACQUITY BEH Amide column (2.1 x 100 mm, 1.7 μM) column and Buffer A contained 0.1% formic acid and 1 mM ammonium acetate and Buffer B contained 95% acetonitrile, 0.1% formic acid, and 1 mM ammonium acetate. The HILIC method was run at a flow rate of 0.3 mL/min, and the gradient began at 90% B and ramped to 20% B from 0.6 min to 3.15 min and held for 2 min. The C18 method consisted of a Thermo Hypersil Gold C18 column (2.1 x 100 mm, 1.7 μM) and Buffer A contained 1 mM ammonium acetate, and Buffer B contained 99% acetonitrile with 1 mM ammonium acetate. The C18 method was run at a flow rate of 0.3 mL/min, and the gradient began at 1% B and 99% A and then ramped to 99% B and 1% A from 0.5 to 1.25 min, followed by a hold until 5 min. The Thermo ID-X mass spectrometer was operated at 120k resolution with a scan range of 85-1275 *m/z*, and the mass detector was set to Orbitrap. Sheath gas was 50, auxiliary gas was 10, and sweep gas was 1. The spray voltage was 3.5 kV for ESI+ and 2.75 kV for ESI-. MS/MS ion dissociation spectra were collected using data-dependent mode and parallel-reaction monitoring (PRM) mode in normalized HCD % was set to 35%. MS/MS was collected at 60k resolution for MS^1^ scans and 30k resolution for MS/MS scans.

## Data availability

The LC-HRMS data used for the head-to-head algorithm comparison and ccRCC biological validation using pathway alterations are deposited in the Metabolomics Workbench and are available as ST005019, ST005054, and ST005111. The R codes and data for reproducing the head-to-head algorithm comparison and external validation are provided in Zenodo (10.5281/zenodo.21574649). The default metabolite reference database (merged from HMDB and KEGG) is provided in **Supplementary Data 1**, the results of head-to-head comparison between different metabolomics annotation algorithms are provided in **Supplementary Data 2**, an in-house Schymanski Level 1 identified metabolite reference library is provided in **Supplementary Data 3**, the MS2Query validation results are provided in **Supplementary Data 4**, the external validation result summaries are provided in **Supplementary Data 5**, the RT reference database for HILIC and C18 chromatography is provided in **Supplementary Data 6**, the metabolite biospecimen concentration reference database is provided in **Supplementary Data 7**, the metabolic precursor-product relationship database is provided in **Supplementary Data 8**, and the metabolite co-transporter relationship database is provided in **Supplementary Data 9**.

## Code availability

MSMICA is an R package available for non-commercial use on GitHub at https://github.com/jamesjiadazhan/MSMICA under the CC BY-NC-ND 4.0 License. User guide is provided at https://jamesjiadazhan.github.io/MSMICA_manual.

## Acknowledgments

This work was supported by a Henry M Jackson Foundation grant (no. 6130 to D.P.J., Y.M.G., M.R.S.), Sequoia Foundation grant (no. 9168-Emory-01 to D.P.J.), a University of California Los Angeles grant (no. 1918SLA19700 to D.P.J., Y.M.G.), and a University of California Los Angeles grant (no. 1935 G LA835 to D.P.J., Y.M.G.). J.D.P. was supported by the Medical Scientist Training Program Grant, T32GM008169. The authors thank Matthew Ryan Smith for the helpful discussion, and Terrance J. Kavanagh, who provided the LC-HRMS untargeted metabolomics datasets from a mouse experiment with multiple tissue samples collected. This research was supported in part by the Intramural Research Program of the National Institutes of Health (NIH). The contributions of the NIH authors are considered Works of the United States Government. The findings and conclusions presented in this paper are those of the authors and do not necessarily reflect the views of the NIH or the U.S. Department of Health and Human Services.

## Author contributions

J.Z. and D.P.J. conceived the project. J.Z. wrote the MSMICA algorithm code. D.P.J., Z.R.J., J.W, and S.T. contributed to coding development. V.T. and J.Z. processed the LC-MS data. J.W., W.J.C., and J.Z. analyzed the LC-MS/MS data. Z.R.J., J.D.P., and J.W. contributed to the figure revision. M.N. provided data access to the 4 studies used for MSMICA parameter development. J.D.P, R.D.C, N.L.P, and V.A.M contributed to data acquisition. J.Z. and D.P.J wrote the paper. All authors provided editorial feedback and revision of the manuscript. All authors approved of the final content in the manuscript.

## Figure legends

**Extended Data Figure 1.**
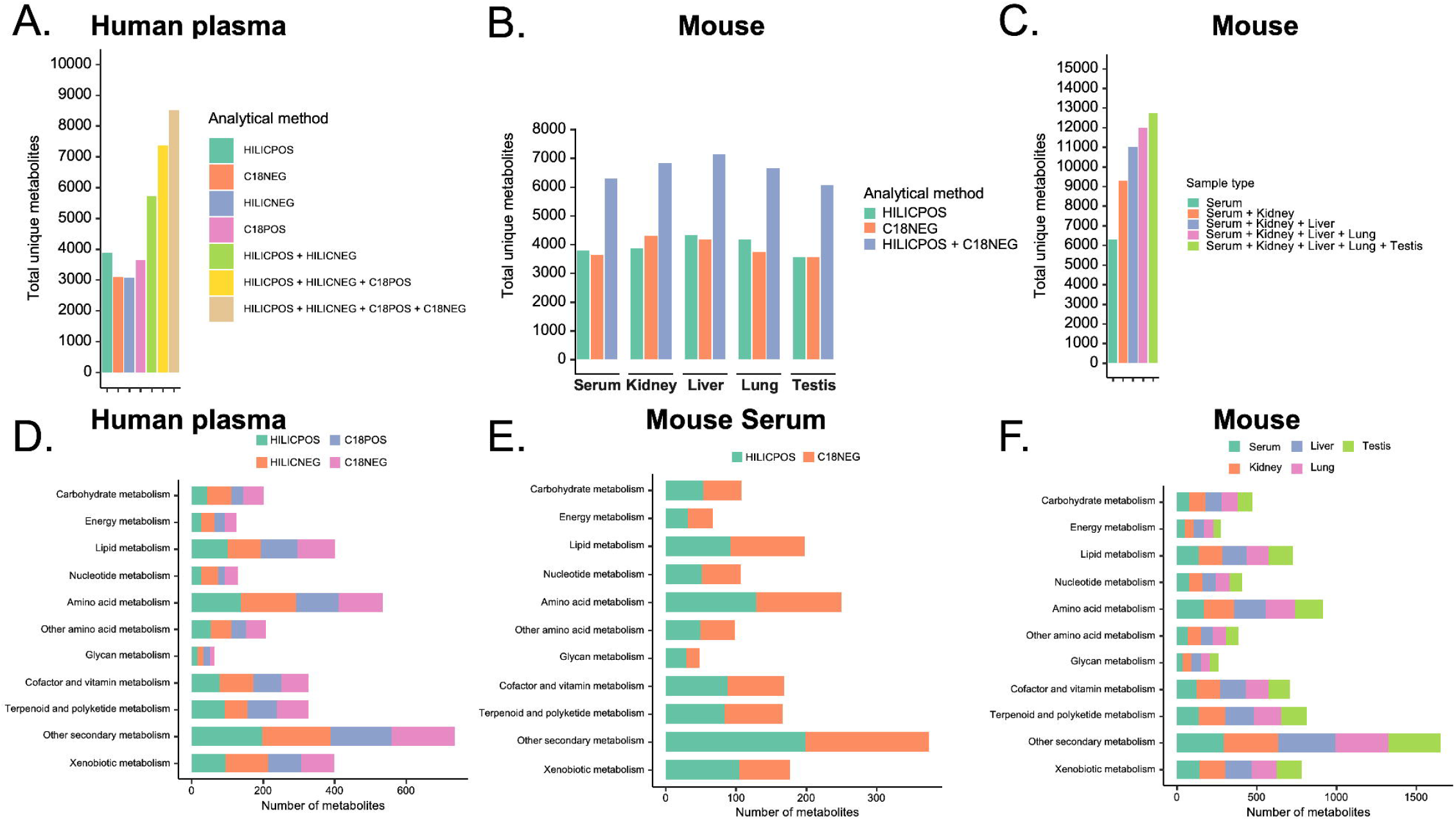
Pathway coverage of MSMICA results. **(A).** KEGG pathway coverages by MSMICA identified results using HILIC+ human plasma data. **(B)**. KEGG pathway coverages by MSMICA identified results using HILIC+, HILIC+, C18+, and C18-human plasma data. Metabolites were deduplicated based on InChIKey. Identity deduplication process was the same for panel C and D. **(C)**. KEGG pathway coverages by MSMICA identified results using HILIC+ and C18-mouse serum data. **(D)**. KEGG pathway coverages by MSMICA identified results using HILIC+ and C18-mouse serum, kidney, liver, lung, and testis data.

**Extended Data Figure 2.**
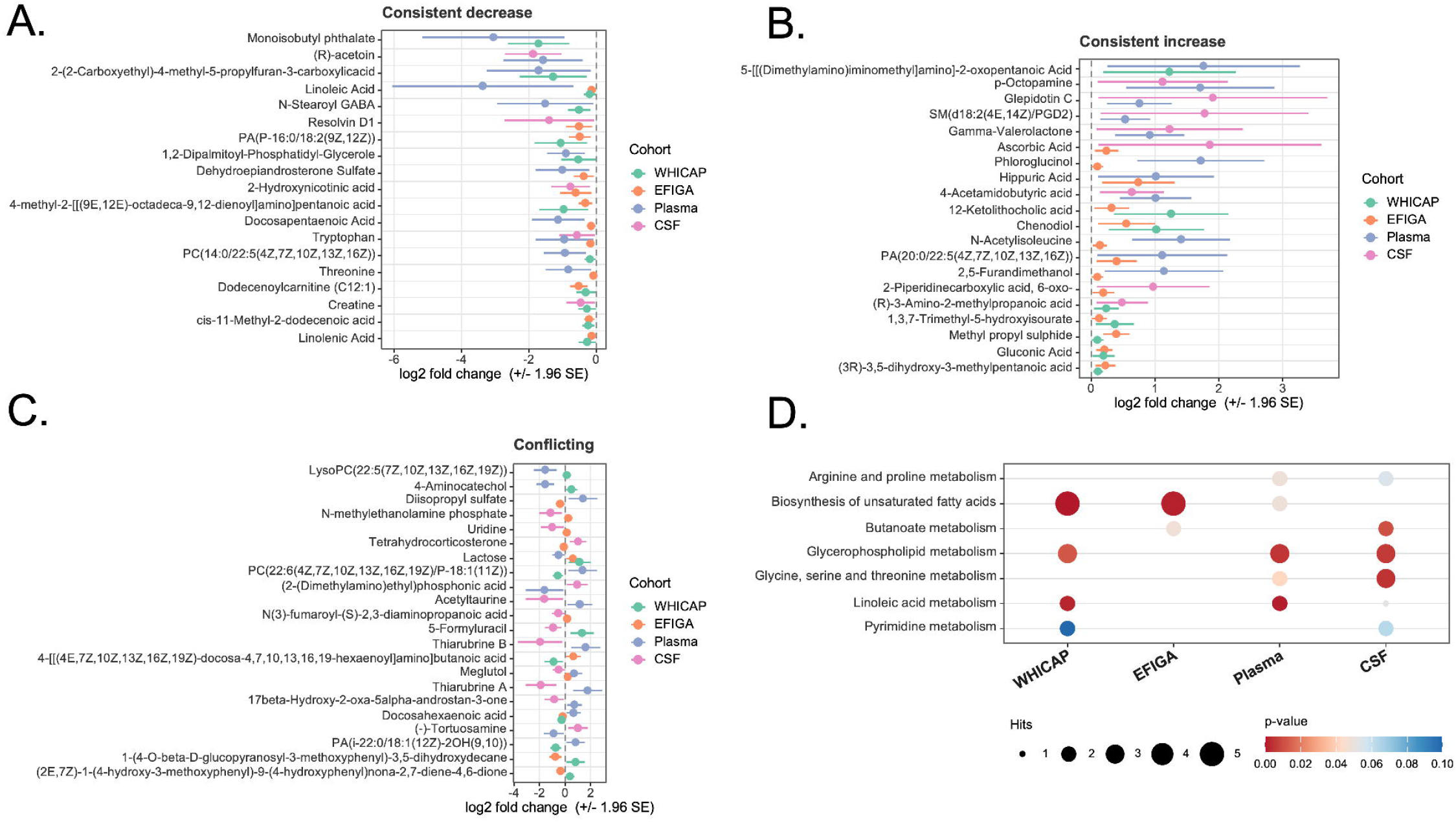
Sensitivity analysis of MSMICA data harmonization analysis. **(A).** Forest plot of metabolites showing consistent decreases across WHICAP, EFIGA, plasma, and CSF datasets. In the WHICAP and EFIGA studies, age and sex were adjusted in the limma analysis. **(B)**. Forest plot of metabolites showing consistent increases across datasets. **(C)**. Forest plot of metabolites with conflicting directions across datasets. **(D)**. Bubble plot summarizing overlapping pathways in at least two datasets.

