## Supplementary Materials for "MSMICA: computational metabolite identification in untargeted metabolomics by integrating MS, retention time, and biological evidence"

**Supplementary information**

### **Supplementary Tables:**

#### **Supplementary table 1**. Characterization of MSMICA workflow

| **Step** | **Description** | **Example data** |
| --- | --- | --- |
| **Input** | Total extracted features | 13805 unique features |
| **1** | m/z matching of features with metabolite adducts without considering isotopic distribution | 171542 metabolite adducts  7511 unique features |
|  | m/z matching of features with metabolite adducts with the most abundant isotopologue | 167174 metabolite adducts  7309 unique features |
| **2** | Parameter estimation using *m/z* uniquely matched metabolites | 1242 metabolite adducts  413 unique features |
| **3** | Clustering of adducts and isotopes using correlations and retention time thresholds | 1917 metabolite adducts  1292 unique features |
| **4** | Local Bayesian optimization per monoisotopic mass | 2796 metabolite adducts  2667 unique features |
| **Output** | Schymanski level 3a results  Schymanski level 3b results  Schymanski level 4 results | 563 metabolite adducts, 561 unique features  2231 metabolite adducts, 2106 unique features  556 metabolite adducts, 512 unique features |

#### **Supplementary table 2**. Category of MSMICA results regarding Schymanski identification level

| **Schymanski identification level** | **MSMICA criteria** |
| --- | --- |
| Level 3a | *m/z* matching  retention time prediction, clustering of adduct and isotopes  biospecimen-specific concentration, precursor-product/transporter correlation |
| Level 3b | *m/z* matching  retention time prediction, clustering of adduct and isotopes |
| Level 4 | *m/z* matching  Secondary adducts or isotopes of level 3a and 3b results |

### **Supplementary Data (in separate downloadable files):**

#### **Supplementary Data 1.** HMDB and KEGG reference compound database

#### **Supplementary Data 2.** Head-to-head comparison between metabolite annotation algorithms

#### **Supplementary Data 3.** In-house Schymanski level 1 identification reference library

#### **Supplementary Data 4.** In-house validation using MS/MS database matching

#### **Supplementary Data 5.** External validation of MSMICA results

#### **Supplementary Data 6.** Retention time reference database for retention time mapping and prediction

#### **Supplementary Data 7.** Metabolite concentration reference database

#### **Supplementary Data 8.** Metabolic precursor-product relationship database

#### **Supplementary Data 9.** Metabolite co-transporter relationship database
